# Assemblatron: An Automated Workflow for High-Throughput Assembly of Big-DNA Libraries

**DOI:** 10.1101/2025.06.30.662359

**Authors:** Antonio Vela Gartner, Luisa F. Quispe, Klaus-Peter Rühmann, J. Andrew Martin, Jordan M. Welker, Mingyu Kenneth Li, Camila Coelho, Yu Zhao, Ran Brosh, Jef D. Boeke

## Abstract

The ability to synthesize 50-kb+ DNA molecules has tremendous potential in the fields of genome engineering, metabolic engineering, and synthetic regulatory genomics. Despite tremendous achievements in these fields, such as the completion of the first synthetic eukaryotic genome, assembling custom big DNAs remains slow, expensive, and laborious. In this work, we present a set of improvements to yeast-based DNA assembly methods that enable medium-to high-throughput big DNA experiments. In particular, we i) developed two easy-to-use vector systems: Jack In the Box (JIB), which deploys a split marker and split centromere strategy to reduce background, and Selection of URS Recombinational Excision (SURE) which exploits URS silencer removal from the *LEU2* marker. These strategies sharply decrease the number of colonies containing an empty vector, greatly reducing the amount of screening required to find correct clones. ii) We improved yeast transformation efficiency, increasing the likelihood that all required DNA segments are co-transformed into the same cell. iii) We developed an automation pipeline for segment mixing through single-colony isolation. We also established three phenotype-driven assays that directly and rapidly report the frequency of correctly assembled large DNA molecules. Harnessing these improvements, we show a high success rate in assembling diverse big DNA libraries by combinatorial assembly and oligo-guided architecture in yeast (OLIGARCHY), a novel method that allows cost-effective reuse of segments for distinct assemblies with reduced effort and cost. Using OLIGARCHY, we designed 96 structural variants of a ∼73kb construct, and a single person was able to correctly assemble 90 of these in 7 days, for a total of 7 Mb of assembled DNA. Finally, we used the SURE vector to greatly improve existing methods to turn oligonucleotides into gene-sized double-stranded DNA segments, efficiently turning 80 overlapping 60-mers into ∼3-kb DNAs directly, without PCR amplification.

## Introduction

The vast majority of molecular biology has been carried out studying one gene at a time, usually with plasmid vectors in the 2-15 kilobase (kb) range, and usually the gene of interest is studied in a “cDNA/transgene” format, lacking its native regulatory and intronic sequences. Big DNAs of 50-kb and up can accommodate substantial gene-flanking regions and can also recapitulate the complexities of alternative splicing so commonly seen in the genes of mammals, plants, and other complex eukaryotes. We typically build big DNA “assemblons” of 50-to 200-kb using the yeast *Saccharomyces cerevisiae* as the microbial engine for assembly^1–3^. These big DNAs are assembled from a set of 20-50 overlapping DNA fragments in the 1-to 5-kb range, sourced either from commercial vendors or by PCR amplification, and a Yeast Assemblon Vector (YAV). Similar methods have been used to assemble whole prokaryotic and eukaryotic chromosomes^4–17^. For these types of projects, it is desirable to increase assembly efficiency to enable the production of much larger numbers of big DNA constructs, and to do so in a library format, in which a set of related but structurally and/or genotypically varying big DNAs can be assembled in parallel. It is also crucial that methodological improvements do not interfere with DNA content—i.e., that designers of big DNAs retain total control over produced sequences. Yeast assembly, powered by homologous recombination (HR), is ideal for this purpose because, when properly applied, it is sequence-agnostic, its fidelity is exceedingly high, and there is thus no need to remove instances of specific sequences like restriction sites or recode open reading frames from the designer sequence of choice. This is in contrast to the Golden Gate method, which is arguably the most efficient and precise in vitro assembly method, but is limited in some contexts due to the need to avoid specific Type IIs restriction enzyme sites in the final assembly^18^.

Here we describe a collection of new technologies underlying the **Assemblatron**, a workflow that powers the yeast-based assembly of many big DNAs in parallel and at unprecedented efficiency and precision, without sacrificing sequence flexibility. Two crucial elements of the workflow are novel ultralow-background vector systems, **Jack in the Box (JIB)** and **Selection of URS recombinational excision (SURE)**, and methodological enhancements of yeast transformation at scale. Multiple steps in this workflow have been automated or are highly amenable to readily available automation platforms, driving costs down.

## Results and Discussion

### 1) Easy-to-use Vector Systems to Select for HR-ready Cells

Yeast DNA assembly typically uses an in vitro linearized vector, and a series of one or more overlapping insert fragments as outlined in **Fig. 1A**. The overlapping insert fragments are recombined precisely by the endogenous yeast homologous recombination machinery into a seamless big DNA “product” (sometimes referred to as passenger DNA or cargo) that can be later studied directly in yeast or delivered to another type of cell such as a mouse embryonic stem cell. It works because yeast interprets the different segments with free ends as double-strand breaks and uses homologous recombination to fix them. Yet, since HR is only active at certain cell cycle stages^19^, undesired pathways like non-homologous end joining (NHEJ) and microhomology-mediated end joining (MMEJ) can lead to mis-assembly, including re-circularized plasmids with no inserts and partial assemblies with missing or incorrectly assembled segments. Screening yeast clones for correct assemblies is one of the most time-consuming steps. To reduce the number of background colonies from re-circularization, a common strategy is using proportionally less vector. However, this also leads to a reduction in colony yield^20^. Given that yeast HR and NHEJ are active in different phases of the cell cycle^19^, a better alternative is to couple plasmid viability to recombination. One way to achieve this is by engineering a selectable marker in the middle of the insert assembly, far from the centromere and ARS sequences^21^, but this approach is not a viable option for some applications like human locus dissection where having selectable marker sequences in the middle of the insert assembly is undesirable^1,2^. A second method involves splitting the selectable marker between 2 overlapping fragments so that it is only functional when reconstructed by HR^22^. While efficient at reducing background, this method typically requires extra steps, thus leading to low adoption. Here, we present the ultralow-background **Jack-In-the-Box (JIB)** and **Selection of URS Recombinational Excision (SURE)** systems as alternatives that select for progeny molecules assembled by HR, yet are as easy to use as regular yeast assembly plasmids.

**Figure 1.**
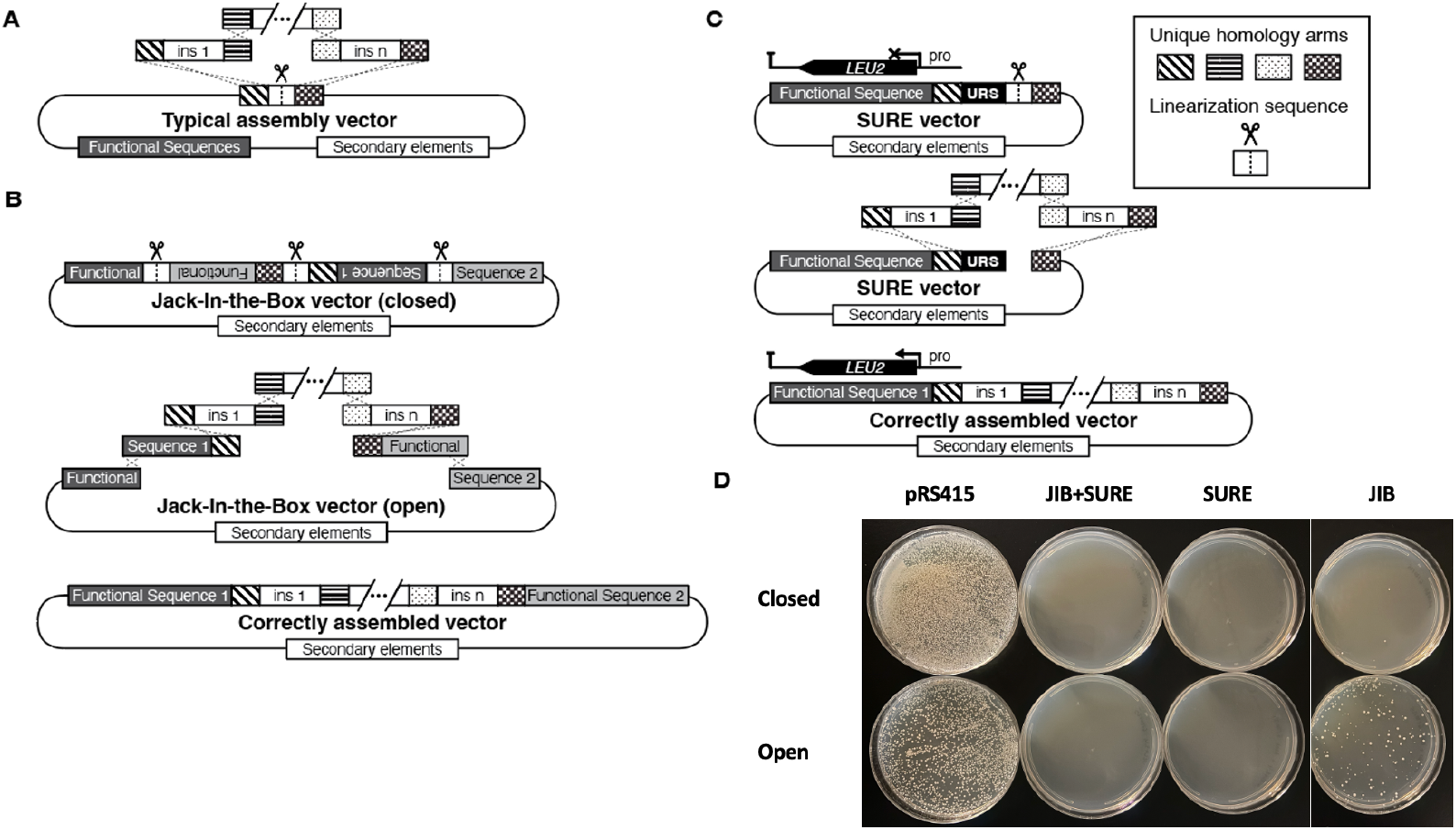
Jack-In-the-Box and Selection of URS Recombinational Excision. **A**. Typical yeast assembly vector. Functional sequences are genes or segments essential to produce a functional vector in yeast. Typically, these would include a selectable marker such as *LEU2*, and a critical element for replication and segregation such as CEN6-ARS1. Secondary elements include genes or segments important for selection and maintenance in *E. coli*. **B**. JIB vector, top shows closed (undigested) configuration; middle shows open configuration following BsaI digestion and indicate the structure of the insert fragments to be assembled; bottom shows end product. **C**. SURE vector. **D**. BY4741 was transformed with 10 ng of pRS415, JIB+SURE, SURE, and JIB vectors transformed without insert fragments in open or closed conformations. All vectors were digested with BsaI.

#### 1.1) Jack-In-the-Box

We define two “functional segments” required for any yeast assembly vector, one of which is a selectable marker, which in our example is the selectable *LEU2* gene which confers the ability to grow in the absence of leucine to a *leu2* mutant yeast, and the centromere segment *CEN6* which in conjunction with the adjacent Autonomously Replicating Sequence (ARS) allows for near single-copy maintenance of circular yeast assembly vectors. The JIB system consists of plasmids that exist in two distinct states, “closed” and “open” (**Fig. 1B**). A closed JIB plasmid can transform

*E. coli* but not yeast since the yeast functional segments, *LEU2* and CEN6, are each broken/split into 2 overlapping but dysfunctional halves: the two segments flanking the cloning site where the insert fragments will ultimately reside contain i) half of the *LEU2* marker and ii) half of the functional centromere *CEN6*, and the third piece contains the remaining functional segment halves plus an *E. coli* shuttle vector. In the closed state, the three segments are cloned in a nonfunctional configuration in a single circular plasmid that is unable to transform yeast cells because neither the *LEU2* gene nor the *CEN* sequences are intact. The open JIB is enabled by BsaI restriction digestion at the 3 linearization sequences (LSs). This produces an open JIB plasmid comprising 3 fragments that overlap at 2 locations and harbor terminal overlaps with the separately designed terminal insert fragments. The closed JIB plasmid can be readily cloned in *E. coli* but yields drastically fewer colonies in yeast than regular vectors like pRS415. The open JIB plasmid generates dramatically fewer colonies than regular vectors when transformed with no inserts (**Fig. 1D**). The JIB vector we have worked with the most, pAVG030, is designed to accommodate big DNA in that it includes a BAC origin allowing for single copy in *E. coli*^23^.

Because the overlapping regions of the broken/split segments in the closed JIB vector are adjacent to each other and separated by the midpoint linearization sequence, they can be easily customized using simple *E. coli* cloning techniques following I-SceI digestion (see **Methods**).

Using the above strategy, we have created the JIB vector with appropriate recombination sites “LVec” and “RVec” used in a combinatorial assembly experiment (pKPR125, **Supplemental Table 1**), as well as a 4-piece JIB vector containing an extra constant piece that gets liberated (*e.g*., a marker cassette 1 sequence to readily enable incorporation of mammalian homology arms for mSwapIN^24^ in plasmid pJMW29).

Yet another type of JIB vector variant we have constructed includes the pUC19 high copy origin rather than BAC origins like pKML070 (JIB with LVec and RVec) and pKML072 (JIB+SURE, see below) which support high copy number in *E. coli;* these will provide higher yields for smaller insert sizes which is helpful for transfer of plasmids to bacteria (**Supplemental Table 1**).

#### 1.2) SURE

A disadvantage of JIB and previous split marker systems is that they increase the number of segments in the assembly, reducing assembly efficiency^1^. To address this, we sought to develop a method to further select for the cloning of any arbitrary insert. Several selection schemes have been described to enhance *in-vitro* cloning methods like TOPO, Gateway or Golden Gate cloning that rely on disrupting or eliminating a negative selectable (counter selectable) marker gene, such as ccdB in bacteria^25^ and TPK2 in yeast^26^, so that only cells with an insert survive. Here, we develop a method of insert selection that depends on active yeast HR. As illustrated in **Fig. 1C**, the recombination event simultaneously inserts the DNA payload while deleting an upstream repressor sequence (URS) that otherwise prevents the expression of a positive selectable marker such as *LEU2*. An advantage of this system is that it only requires a small but potent regulatory sequence, thus keeping plasmids compact, while not adding any further selection marker or drug. This is unlike traditional methods, which suffer from background caused by frameshifts in the negative selectable marker or by plasmid re-circularization after marker removal. We call this system Selection of URS Recombinational Excision, or SURE. Our SURE plasmid was made by appending the *INO2* URS sequence^27^ to the core *LEU2* promoter (*i.e*., the sequences distal of the known Leu3 upstream activator sequences) found in pRS415. We confirmed that both circular and linearized SURE vectors do not yield Leu^+^ colonies (**Fig. 1D**). The core (minimal) *LEU2* promoter works by itself, but yeast colonies grow faster when the 22bp Leu3 UAS sequence is restored as part of the insert (**Supplemental Figure 1**). We also made a JIB+SURE vector by modifying a JIB vector and showed it has the same low background (**Fig. 1D**).

### 2) Phenotype-based assays for assessing assembly validity without sequencing

### 2.1) The YAV8 system

Big DNA yeast assembly is an inordinately powerful but also complex process that relies on the successful completion of dozens of homologous DNA recombination events that seamlessly stitch together the 30-50 overlapping “insert” segments with a suitable vector. A typical assembly project consists of about 33 insert segments overlapping each other by 60-200 bp and cloned into a Yeast Assembly Vector^1–3,5,21^. Assembly verification involves isolating the cloned DNA and subjecting it to DNA structural analyses such as PCR genotyping, digestion with restriction enzymes, and/or DNA sequencing. While these are well-developed methods, comprehensively analyzing tens to hundreds of such clones is time-consuming, expensive, and error-prone.

Implementing a fast, high-throughput phenotypic assay to verify assemblies, rather than relying on sequencing, would dramatically streamline and scale the optimization of the assembly process. We therefore developed such an assay based on a set of seven different large auxotrophic marker genes from seven distinct biosynthetic pathways for amino acids and nucleotides, each ranging in size from 2400 to 6645 bp (**Supplemental Table 2**). We then flanked these sequences with inert stuffer sequences from the yeast *Schizosaccharomyces pombe* genome to attain a total insert+vector size of 98,246 bp, of which about a third consists of selectable marker open reading frames (ORFs) or functional vector sequences (**Supplemental Figure 2**). With this design, when the insert is segmented into segments of ∼3000 bp, about a third of the overlap regions lie within selectable regions, and thus misassemblies are readily identified by the loss of function of one or more of the 7 marker genes *ADE6, ARG56, ARO1, HIS4, LYS2, MET5*, and *URA2* that make up the “insert” portion of YAV8. The 8^th^ selectable marker, *LEU2*, is part of the vector backbone and is used for selection during transformation. The additional element of the YAV8 system we constructed is a suitable yeast strain (yHB035) with deletions of the 8 auxotrophic marker genes (**Supplemental Table 3**).

Thus, following an assembly transformation of the vector backbone segment(s) and the YAV8 insert segments, the transformants were selected on plates lacking leucine, selecting only for the vector backbone. These colonies are subsequently replica-plated (or otherwise tested for growth) on SC-8 plates lacking all 8 nutrients. Successful growth indicates that a substantial proportion of the YAV has been assembled correctly, serving as a proxy for proper assembly in the relevant context, namely *de novo* assembly from 30-35 overlapping fragments (**Fig. 2**).

**Figure 2.**
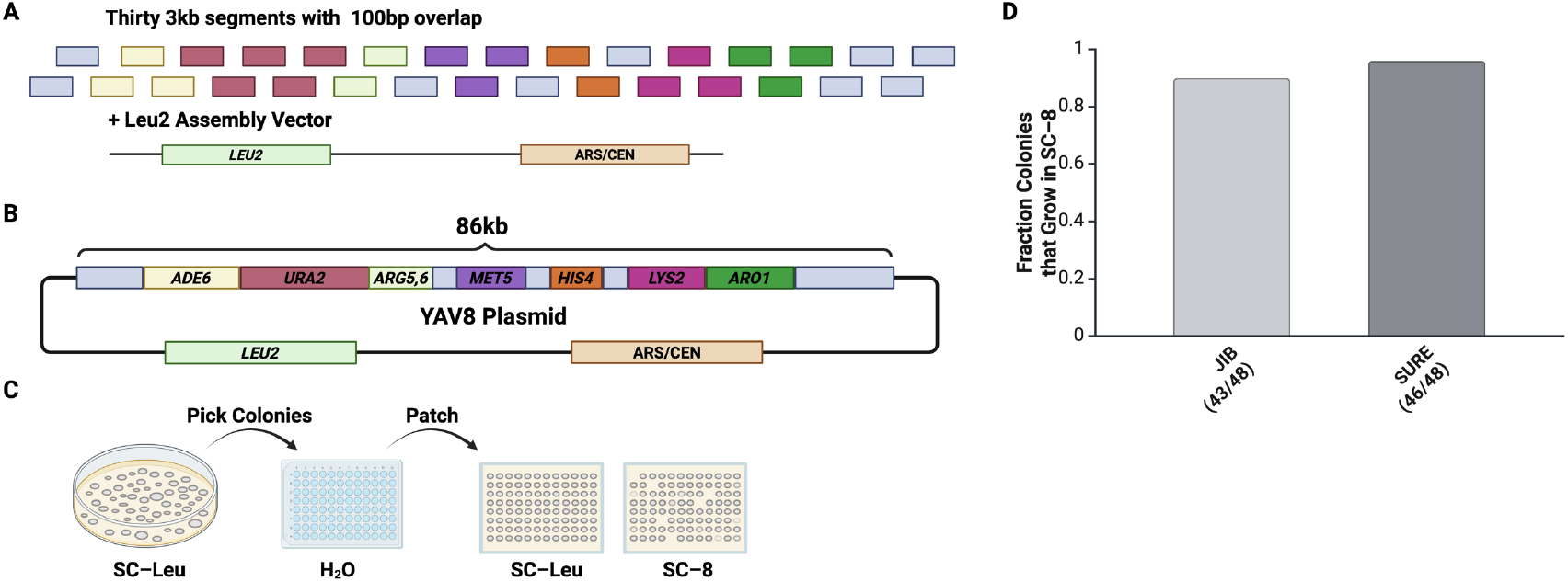
Using the YAV8 vector system to test JIB and SURE. **A**. YAV8 assembly using 30 ∼3-kb amplicons and a *LEU2* Vector. **B**. A successfully assembled YAV8 plasmid contains the full 86-kb insert in a *LEU2* assembly vector. **C**. Selection scheme to identify correctly assembled YAV8. The ratio of fully auxotrophic colonies to Leu2+ colonies determines assembly precision. **D**. JIB and SURE YAV8 assembly precision.

#### 2.2) YAV8 system as a performance assay for JIB and SURE vectors

We used the YAV8 system to show that the JIB and SURE vectors yield high assembly accuracy (**Fig. 2**). In short, we amplified 30 ∼3-kb segments, overlapping by 100 bp from a plasmid containing the 86-kb YAV8 region, mixed them at equimolar ratios, and transformed with the indicated assembly vectors. Note that segment #1 was amplified to match each vector’s end and add a Leu3 UAS when needed, while segment #30 was amplified to have a common homology to all our vectors. Transformants were selected on SC–Leu and replica plated onto SC–8. Both JIB and SURE vectors had assembly accuracy of > 90% (**Fig. 1D**). When the YAV8 insert is assembled into the SURE vector, it forms pAVG058; when using the JIB vector, it forms pAVG059. We picked two colonies from the SC–8 patches for experiments and sequenced the assemblies by nanopore whole genome sequencing; for pAVG058 one was sequenced perfectly and the other one had a transition mutation (probably from PCR), for pAVG059 both were sequence-perfect.

#### 2.3) Improved multiplexed oligonucleotide assembly of gene-sized DNA constructs

Given the outstanding performance of these vectors with multiple double-stranded (ds) DNA segments, we wondered whether a similar success could be achieved with single-stranded (ss) DNA parts. In 2009, Gibson showed that it was possible to build a 1.2-kb DNA fragment from 38 fully overlapping 60-mer oligonucleotides (oligos), obtaining complete assemblies 66% of the time^28^. We used our SURE vector to build a 2.5-kb DNA composed of *URA3* (the same DNA sequence built by Gibson) plus *SpHis-5* from 81 fully overlapping oligos, obtaining complete functional assemblies that grow on medium lacking uracil and histidine 80% of the time (**Fig. 3**).

**Figure 3.**
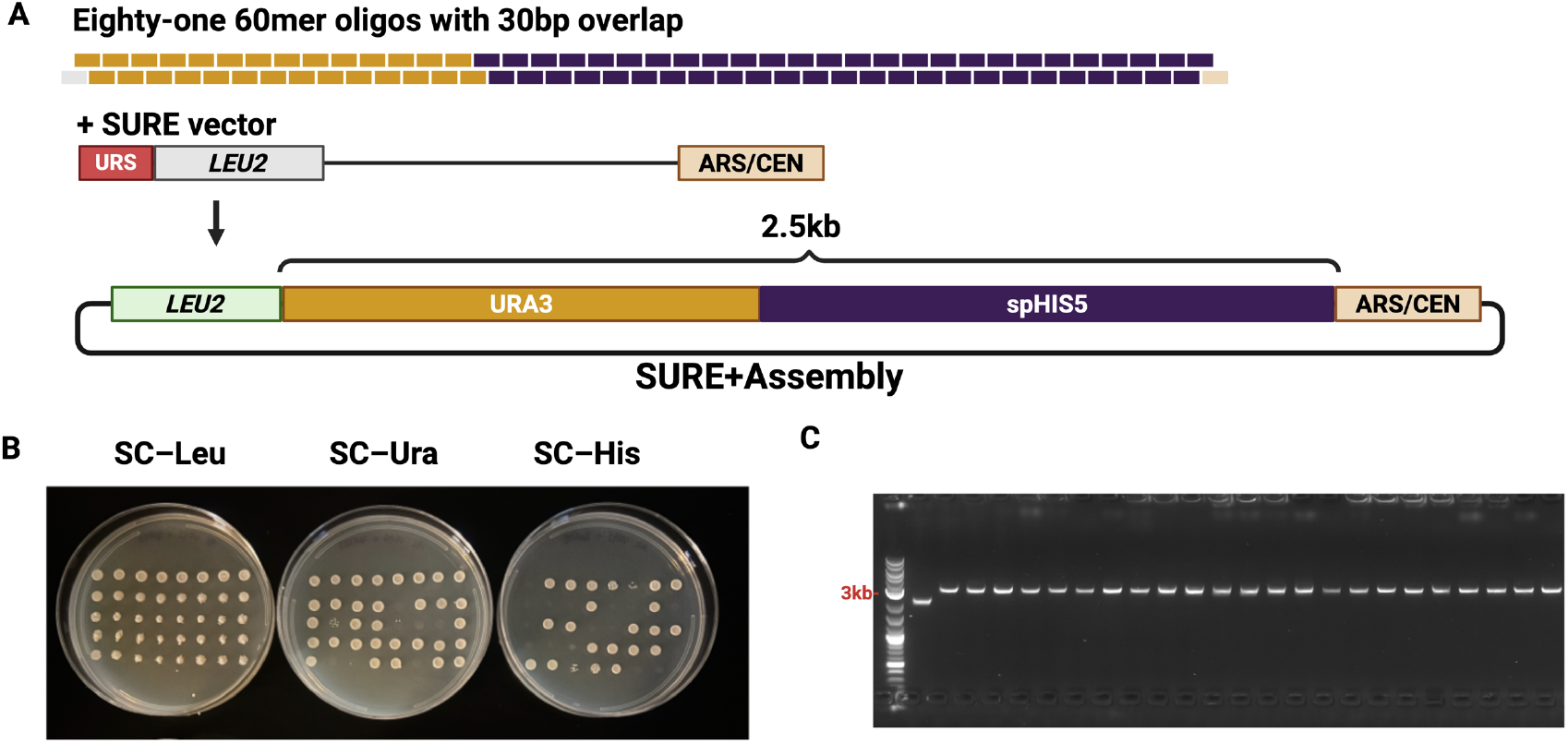
High fidelity assembly of a 2.5kb DNA from 60mer oligonucleotides using a SURE vector. **A**. Assembly diagram. Gold (*URA3*) and black bars (*spHis-5*) represent perfectly overlapping 60-mer oligos. The grey oligo at the left end overlaps LEU2 and tan oligo at the right end overlaps ARS-CEN. SURE vector pAV052 is indicated. **B**. Replica-plating shows that most of the Leu^+^ colonies had intact *URA3* and *SpHis-5* sequences in them. **C**. Gel showing that all colonies tested contained an amplicon consistent with correct assembly; many of these were either Ura^−^, His^−^ or both, presumably as a consequence of the high innate error frequency of the provided oligonucleotides.

### 2.4) Pooled assembly of a phenotype-based Big DNA library enabled by the JIB vector system

We tested the JIB vector system in the context of combinatorial assembly^29^ for the construction of high complexity big DNA libraries, an application where the use of an ultralow-background vector system can be extremely helpful in maximizing both library complexity and yield of correctly assembled DNA constructs. To model this approach and rapidly generate a combinatorial biosynthetic pathway library, we applied our strategy to shuffle a set of distinct DNA fragments, forming a yeast big DNA library capable of producing colonies directly distinguishable by pigmentation and/or fluorescence phenotypes. Individual transcription units (TUs), each consisting of a native yeast promoter, a codon-optimized gene, and a native yeast terminator, were grouped into six pre-defined “slots” and flanked by 100-bp homology arms specific to their assigned slot (**Fig. 4**). Yeast transformation of an equimolar pool of TU fragments, together with open JIB vector backbone fragments, enabled formation of a complex library, designed to allow only one TU per slot to be incorporated.

**Figure 4.**
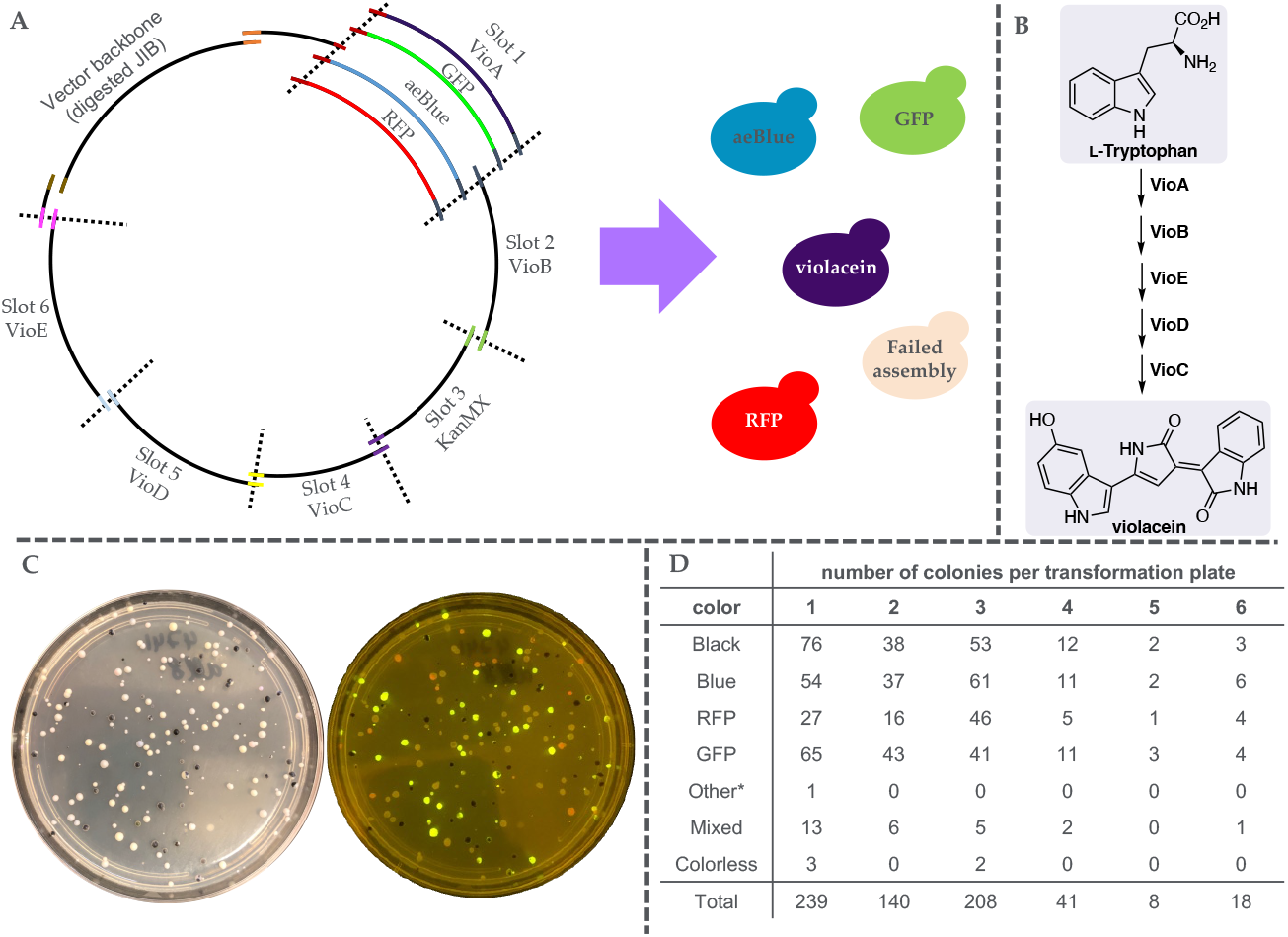
Color assay to test combinatorial yeast assembly. **A**. Schematic overview: Deploying the JIB vector and nine transcription units (TUs) grouped into six assembly “slots”. Each TU is flanked by two 100 bp terminal homology arms (colored segments). Yeast transformation using equimolar quantities of TU fragments is expected to produce dark purple/black (violacein), blue, red, and green fluorescent colonies in roughly equal proportions. **B**. Violacein biosynthesis pathway. **C**. Representative agar plate showing yeast colonies of all colors from a combinatorial assembly transformation, visualized under ambient light irradiation (left) or fluorescence (right). **D**. Table summarizes number of colored colonies from six representative transformations, demonstrating a well-balanced distribution of genes among the assembled plasmids. Less than 1% of colonies were colorless. *1 colony was misassembled and produced a green visible color known to be due to partial activity of the *vio* pathway lacking *vioD*.

Building on previous work, we designed a colorful proof-of-concept experiment consisting of six slots encoding the full biosynthetic pathway for the bacterial pigment violacein (VioA–E), along with an additional selectable marker (KanMX)^30^. To visually track combinatorial outcomes, we supplemented the assay with three synthetic DNA fragments encoding GFP, RFP, and the chromoprotein aeBlue, each bearing the same homology arms as VioA and thus competing for integration as the first fragment. When transformed into yeast using equimolar fragment pools, this system yielded colonies with diverse pigmentation and fluorescence profiles, reflecting successful and unbiased assembly across all six slots and the vector (**Fig. 4**). Essentially 100% of the colonies that formed were either purplish-black (indicating intact violacein pathway and *vioA* gene in slot 1), light blue in color, or fluorescent. A minority of colonies showed a multicolored or color + fluorescent phenotype, presumably reflecting polyclonal or mixed colonies and these were not studied further. These results confirm that the JIB vector system has an ultralow background and that this library assembly strategy supports an efficient, scalable, and balanced construction of combinatorial plasmid libraries, providing a robust foundation for pathway shuffling and downstream screening. An example of how such a library could be used to screen for pathway composition and/or architectures that enable identification of the optimal pathway composition/architecture from a very broad set of starting DNA fragments is shown in **Supplemental Fig. 3**.

### 3) Automating Yeast Assembly: the Assemblatron

To date, true end-to-end automation of yeast DNA assembly remains elusive: the most advanced system described is only semi-automated and suffers from several practical limitations^20^. It uses bulky deep-well plates that consume valuable space and must be manually shuttled between the liquid handling robot, an incubator, and a centrifuge. In that method, only a few steps were automated: (1) mixing of a subset of DNA segments, (2) mixing of the cells with PEG, carrier ssDNA, and DMSO, and (3) transformation patching, leaving all intermediate transfers to the incubator and centrifuge to be performed by hand. Another study describes automated yeast transformation of small constructs made by Golden Gate. However, the transformation efficiency is low, the incubation time is four times longer, and getting single colonies depends on tuning the amount of yeast pipetted during the transformation step^31^.

Here we demonstrate the first fully automated platform for large DNA assembly, integrating all steps from segment mixing to single colony formation, and a downstream automated single colony purification method (Fig. 5), deployed on an Opentrons Flex robotic liquid handler. The platform’s efficiency in producing big DNA depends on the ultralow background conferred by the JIB or JIB/SURE systems. We refer to the workflow underlying this automated big DNA system as the Assemblatron.

**Figure 5.**
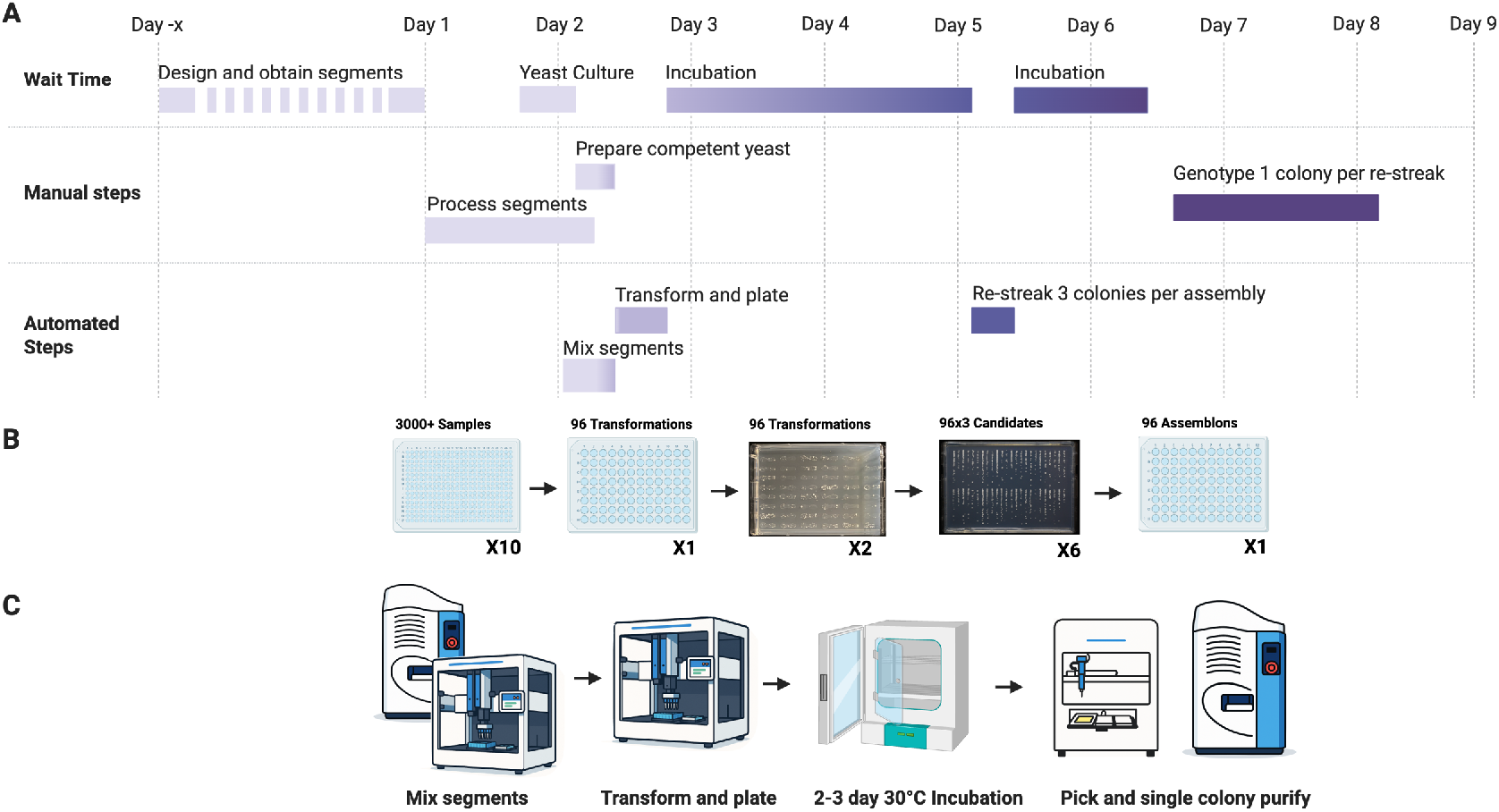
Assemblatron overview. **A**. Typical timeline for producing ∼1 Mb of DNA starting with 3000-mer insert fragments and JIB or SURE vectors. **B**. Overview of Assemblatron sample types. **C**. Automation devices used in the Assemblatron.

#### 3.1) Miniaturization of yeast transformation increases transformation efficiency and enables automation

To achieve this, the first innovation required was miniaturization of yeast transformation, allowing it to be performed efficiently in 96- and 384-well plates. We first attempted to transform with only 30 µL of transformation mix (representing 1/10 of the usual amount in our standard transformation workflow) but this yielded a transformation efficiency of just 10^5^ CFU/µg (1-2 orders of magnitude lower than can be obtained with larger volumes in test tubes). We reasoned that this was due to using fewer yeast cells. To address this, we modified the proportion of reagents in the transformation to contain the same number of yeast cells as a standard 300 µL transformation but in a total volume of 30 µL. To our surprise, this not only recovered, but dramatically increased efficiency from the expected 10^6^ CFU/µg to about 10^7^ CFU/µg (**Supplemental Fig. 4**). We confirmed these results in both 96- and 384-well formats using only 15 µL transformation volume. We conclude from these and other experiments (see **Methods** section) that by concentrating the cells 10-fold we increase the frequency of random collision between transforming DNA and cells and that this drives much higher transformation efficiency.

A key advantage of using small-well plates is that the yeast can be easily incubated in an in-deck thermocycler or heat block, allowing for excellent heat transfer and convenient reagent mixing, facilitating automation of multiple transformation steps by a liquid handling robot. A detailed protocol is described in Methods.

#### 3.2) Automated plating of yeast transformations with robust single-colony formation

After this stage, the transformants could be plated by hand, but using individual petri dishes for 384 transformations would be wasteful and slow. As such, we devised a novel way to plate up to 48 transformation samples in a rectangular agarose plate (such as a Nunc OmniTray) and, importantly, obtain single colonies with very high reproducibility. After the transformation step, an automated 8-channel pipette aspirates 15 µL of 5% PEG and 5 mM CaCl solution and mixes it into eight transformations at a time. Using the same tips, it then aspirates 15 µL of the mixture and sequentially dispenses 6 droplets of 2.5 µL onto a selective plate in such a way that the droplets form a streak with mechanical dilution along the long axis of the resultant streak of cells. In total, six columns of eight streaks fit comfortably, meaning that 384 samples can fit in 8 plates (**Fig. 5B**). After plating, the selective (SC–Leu) agar plates are incubated at 30°C for three days to allow colony formation.

#### 3.2) Using an acoustic liquid handler for fast segment mixing and single-colony purification

The above protocol works very well but is wasteful in the sense that each segment uses a tip every time it is transferred to a sample. Thus we employed an acoustic liquid handler, which has several other advantages, including accuracy, automation, scale, reliability, as a preferred alternative device for mixing segments^31–33^.

An important step in quality control of yeast DNA assembly is single colony purification, which ensures that colonies used have a single assembly genotype. However, performing single colony purification is slow and difficult to scale, making it a bottleneck when one is picking e.g. 4 candidate colonies each for 96 or more assemblies. We developed a method based on an acoustic liquid handler to quickly perform yeast single-colony purification. After yeast colonies are picked by hand or using a colony picker into 100 µL of water, 30 µL of cell suspension is transferred into a 384-well plate designed for the LabCyte Echo acoustic droplet ejection robot. Using this device, 2.5 nanoliter droplets are sprayed onto a streak at high density at one end of the streak and at steadily decreasing density across the long axis of the streak (**Fig. 5B**, see **Methods** for details).

### 4) Oligo-Guided Architecture in Yeast (OLIGARCHY)

It is unlikely that many labs can afford the starting material required to make 96 different 100-kb assemblies. Even at the lowest current per-base cost available for cloned and sequenced gene-sized DNA we are aware of (9 cents/base), the cost would be >$600,000 for the required starting 3000-mers. A more realistic approach would be to reuse several segments in many variant assemblies^34,35^. However, this often involves either having a predefined set of overlaps or using PCR to adapt segments one by one to have the right overlaps. The former is not ideal for assemblies that require specific sequences, and the latter is very time-consuming. As an alternative, we present Oligo-Guided Architecture in Yeast (OLIGARCHY), a simple way to define the order and direction in which DNA segments are joined using oligonucleotide library-derived linkers (**Figure 6**). In OLIGARCHY, a subset or all segment junctions are not the results of the recombination of segment-segment overlapping sequences but the result of two recombination events defined by small dsDNA linkers. In theory, these linkers can be derived in many formats, but the fastest and cheapest way is to order a large oligonucleotide library in which sub-pools can be selectively amplified to produce the specific linkers required for a specific architecture. This is very similar to existing protocols used to make gene-sized DNAs from oligo libraries^36^.

**Figure 6.**
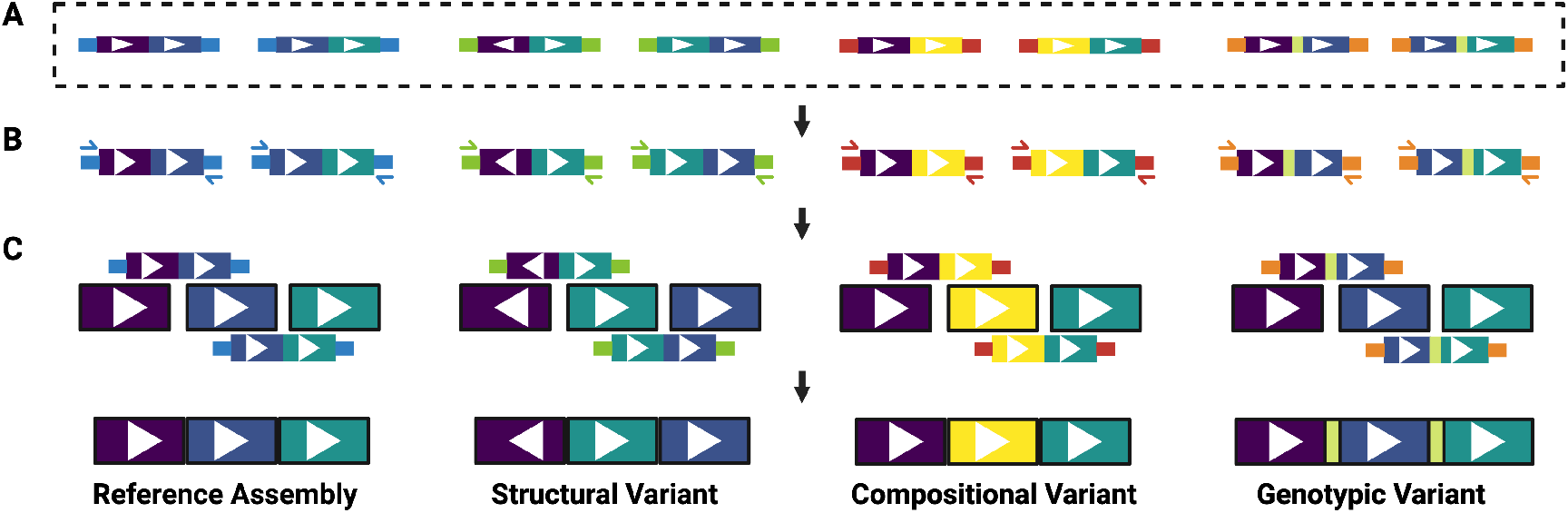
Oligo Guided Architectures in Yeast (OLIGARCHY) overview. Simple diagram of an OLIGARCHY library generated from four different segments joined in groups of three, each by groups of two linkers. **A**. An oligonucleotide library containing all the precursors for the linkers. **B**. Linker groups are extracted from the synthetic oligo library by PCR using primers specific to each group. In the diagram, the primer binding sites sequences are represented by the narrow lines flanking each linker. **C**. Example OLIGARCHY variants. Following PCR amplification, each linker group is transformed into yeast with the appropriate set of segments to generate its variant. Compared to a reference assembly: the structural variant has segment 1 in the opposite direction and segments 2 and 3 in different positions, the compositional variation has a different segment in the second position, and the genotypic variant includes extra sequences brought in by the linkers at the junctions.

#### 4.1) One-step automated assembly of Big DNA structural variants of YAV8

To test this paradigm, we designed 96 structural variants with all the segments from the 86-kb YAV8 region, except omitting segments 2, 14, 18, 21, 28, 30, 31, and 32, which do not carry any essential elements required for the function of the 7 auxotrophic marker genes. These variants are formed by five different sub-assemblies of sizes ranging from 2.9-kb to 31.8-kb joined in different orders and orientations using selectively amplified sub-pools of six linkers (**Figure 7**). As such, each assembly consists of the SURE vector, twenty-three ∼3000-mers, and six linkers with either 134-bp or 50-bp overlaps to the appropriate sub-assemblies or vector. Notably, the first linker also contains the Leu3 UAS sequence between the overlap with the SURE vector and the overlap with the first sub-assembly. This was done purposely to demonstrate that OLIGARCHY can also be used to introduce further designer variability by fitting specific sequences to be added to final constructs inside the linkers.

**Figure 7.**
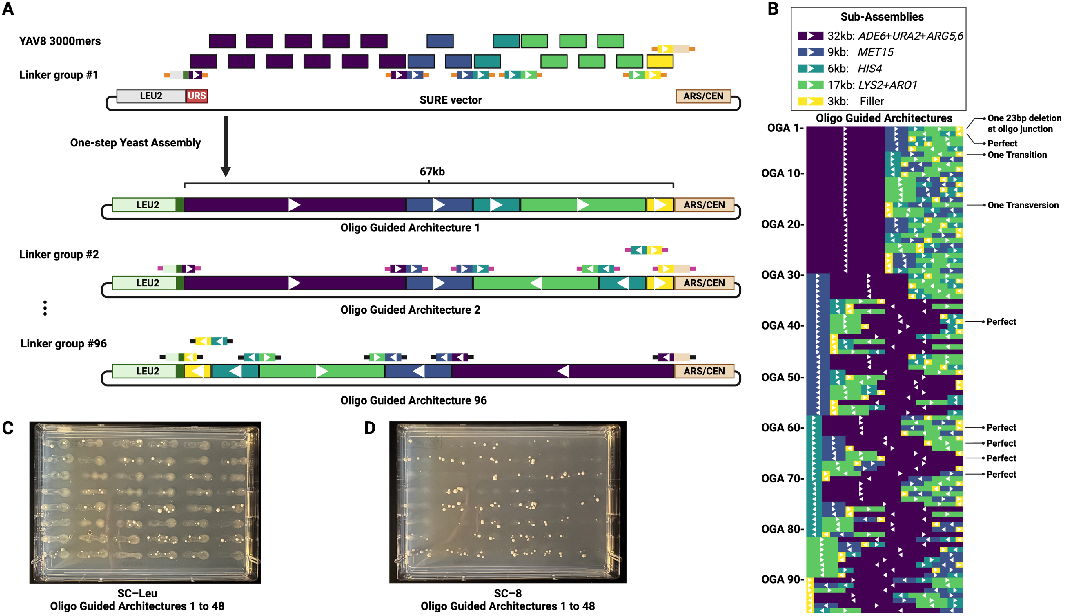
Oligo Guided Architectures in Yeast (OLIGARCHY). **A**. Oligo Guided Architectures (OGAs) to generate structural variants of YAV8. Each structural variant is composed of 5 sub-assemblies linked in different order and directionality. Specific primers are used to amplify linker groups from an oligonucleotide pool; the primer binding sequences are represented by the narrow lines flanking each linker. The left-most linker, which recombines with the SURE vector, introduces a Leu3 UAS (dark green). White triangles are used to signal orientation relative to reference OLIGARCHY architecture #1. Each assembly consists of 23 segments of about 3-kb and 6 linkers. Linkers of 300bp and 132bp were tested, with similar success rates. **B**. Representation of the 96 designed architectures. The assembly of nine colonies that grew in SC–8 was sequenced by Oxford Nanopore sequencing of whole genome preps, the results are annotated to the right of the architecture. **C**. SC–Leu plates of the first 48 OGAs using the 132bp linkers plated from the automated Assemblatron pipeline. **D**. SC–8 plates of the same samples.

Each of the 96 assemblies consisted of an equimolar mixture of the twenty-three YAV8 segments and a custom oligonucleotide pool unique to each assembly. Using our automated pipeline, we mixed all required 3000-mer segments, independently amplified each linker group, and transformed all 96 assemblies. Half of the volume of the transformations was plated on SC–Leu and the other half on SC–8.

We performed the experiment with 134-bp overlap linkers twice; the first time, 95/96 oligo guided architectures (OGAs) had colonies in SC–Leu, with OGA 17 missing. Of these, 90/96 (94%) architectures had colonies in SC–8. In the second repeat, 93/96 assemblies had colonies in SC– Leu, and in this set, OGA 17 was also missing. For the 93 that had colonies, we picked one colony per assembly and patched them to SC–8, 66 of these were positive (i.e., 71%).

For the experiment with oligos with just 50-bp overlaps, 86/96 architectures had colonies in SC– Leu and 81/96 in SC–8. However, four OGAs that did not have colonies in SC–Leu had one or two in SC–8, so that colonies were obtained for 90/96 or 94% of OGAs. Importantly, OGA 34 was among the missing ones. We did observe that with the 50-bp overlap linkers, we had about three times fewer total colonies per transformation. One colony from each architecture in the SC–Leu plates was picked and patched in SC–8, 32 were positive (i.e., 37%). Taken together, these results would suggest that linkers with 50bp overlaps were slightly less effective at assembly, which is not surprising.

## Methods

### Design of JIB Plasmids

The Jack-In-the-Box in the box vector used in this paper, pAVG030 was designed by splitting *LEU2* roughly in the middle of the ORF leaving 100bp of overlap homology. Similarly, *CEN6* (centromere from chromosome 6) was split such that one half will contains CDEI and CDEII and the other contain CDEII and CDEIII, with an overlap of 93 bases. Individual parts of with appropriate homologies were bought and cloned into an inducible bacterial backbone. The JIB parts are cloned separated by linearization sequences (LS). The LS’s were designed to have two divergently oriented BsaI sites that leave incompatible overhangs, separated by an EcoRV site.

### Modification of JIB plasmids

If specific sequences are desired as insert termini, e.g. as homology arms intended for use after excision from the fully assembled vector plus insert, the “cloning site” of JIB vectors can be easily modified by digesting with I-SceI (there are two cut sites that flank the midpoint linearization sequence. Subsequent Gibson assembly^37^ can be used to insert any DNA of interest at the insert cloning site. Importantly, the insert must restore the mLS that is lost upon I-SceI digestion. One limitation to this design is that incomplete digestion will produce plasmids for which the mLS remains uncut, the resulting fragment pieces can recombine with the main vector piece which regenerates a fully functional plasmid and produces low background if linearization digestion is incomplete.

### Design of SURE plasmids

The SURE plasmid was derived from a “previously assembled JIB” (pAVG044) plasmid (this is a partially digested JIB plasmid that was allowed to recombine in yeast to form a functional plasmid without any insert fragments). pAVG044 was digested with BstEII and I-SceI (excising the *LEU2* promoter plus part of the *LEU2* ORF) and a set of 3 gBlocks were cloned into it using Gibson assembly. All of these gBlocks restored the *LEU2* ORF at one end and the LS sequence on the other, one added the full promoter including the Leu3 UAS + 18bp, another added the core promoter lacking the Leu3 UAS, and the third added the core promoter but with the added *INO2*-derived URS1 sequence (ATCCATGCGGAGGCCAAGTATGCGCTT-CGGCGGCTAAATGCGG)^27^. The latter plasmid, pAVG052 is the SURE vector. All three vectors were sequence-verified and transformed into yeast, upon selection for *LEU2* function, the SURE vector yielded no colonies, the other two produced colonies.

### Design of JIB+SURE Plasmid

The exact approach and gBlock used to modify pAVG044 was used to modify JIB1.2 (pAVG030) to yield the JIB+SURE vector (pAVG054).

### JIB and SURE vector linearization

Plasmid DNAs were mini-prepped (Zippy miniprep, Zymo Research, Irvine CA) from *E. coli* strain Epi300, treated with Induction Solution (Lucigen Inc., Middleton WI) or 0.01% arabinose, and digested at a concentration of 5fmol/µL (about 30ng/uL) using 6.6 units of BsaI per µg of vector overnight, and then heat inactivating the enzyme at 80°C for 20min. No gel purification was performed, assuring equimolar production of the three open JIB components. 5 fmol of plasmid vector digest was added to each yeast transformation.

### YAV8 Experiments

The sequence of the original YAV8 vector, pCTC200, is shown in **Supplemental Figure 2**. The YAV8 segments used in these studies were designed using a simple algorithm that iterated primer3^38^ to pick a set of segments of about 3kb that ended with primers with a melting temperature close to 55°C. The segments were amplified using Q5 polymerase or Takara PrimeStar polymerase. The concentration of each segment was measured using gel electrophoresis standards and an Echo robot was used to transfer sufficient volume to specify 8 fmol of each segment per assembly. All transformations were done using miniaturized transformation (see below) either by hand or with the assistance of automation. To test for assembly, colonies grown in SC–Leu were either replica plated to SC–8 or picked in 20µL water (either by hand or with a Hudson RapidPick SP, Hudson Robotics, Springfield NJ) and 5µL were patched into SC–8 (either by hand or using an Opentrons FLEX liquid handling robot).

### Single-stranded short oligonucleotide-based assembly

The oligos used for the *URA3*+*spHIS5* assembly were designed by hand and consisted of 60mers with 30bp full overlaps that tiled the sequence on both strands. Some “padding” was added so that the sequence was an even multiple of 60bp. Additional oligos were added to have homology to the SURE vector and add the Leu3 UAS sequence. The sequences of the oligonucleotides are provided in **Supplemental Table 4**. The oligos were ordered at a concentration of 100µM and combined (by hand or using an Echo as above) to have a concentration of 1µM each, then they were boiled and slowly cooled down to anneal them. The annealed oligos were transformed with 5fmol of predigested SURE vector.

### Standard Lithium acetate yeast transformation conditions

An overnight culture of a yeast strain of choice was grown overnight in appropriate media (e.g. YPD or appropriate SC-dropout medium), and a subculture was made appropriate to the number of transformations to be done. For example for a 12 DNA transformation experiment, a 60 mL culture diluted from an overnight culture to an A_600_ of 0.15 and grown to an A_600_ of 0.4-0.6 (about 3.5 hours for BY4741 or yAVG01). Cells were pelleted at 1500 g for 3 min, washed with an equal volume of sterile water, and finally with a 0.1 M LiOAc solution, and the cell pellet was resuspended in the residual supernatant after discarding bulk LiOAc solution. The resuspended cells were then concentrated by a quick spin in a microfuge and resuspended in 0.6 mL of 0.1 M LiOAc. A mixture of 44% PEG-3350 (2880 µL), 1 M LiOAc (432 µL), 10 mg/mL boiled herring sperm DNA (300 µL per sample) was prepared and mixed with the cells. 300 µL of that mixture was combined with each DNA to be transformed (the DNA added was included in a maximum of 50 µL volume). Final concentrations are approximately as follows: PEG 30%, LiOAc 0.12 M, boiled herring sperm DNA 0.7 mg/mL, cell equivalent A_600_ 5.7–8.6.

### Miniaturized Transformation

An overnight culture of a yeast strain of choice was grown overnight in appropriate media (e.g. YPD or appropriate SC-dropout medium), and a subculture was made appropriate to the number of transformations to be done. For example, for a 12 DNA transformation experiment, a 30 mL culture diluted from an overnight culture to an A_600_ of 0.15 and grown to an A_600_ of 0.4-0.6 (about 3.5 hours for BY4741 or yAVG01). Cells were washed as above except the final pellet was resuspended in a mixture of 44% PEG (144 µL per sample), 1 M LiOAc (21.6 µL per sample), 10 mg/mL herring sperm ssDNA (15 µL per sample). The DNA sample to be transformed consists of 5-8 fmols per insert fragment and vector fragments. When mini-transformation is done is multiwell plates, 15 µL of the cell mixture was added to each well, together with DNA samples in a volume of at most 2.5 µL. Note the changes in final concentrations: PEG 35 %, LiOAc 0.12 M, boiled herring sperm DNA 0.83 mg/mL, cell equivalent A_600_ 66.7–100.

### Automated Yeast Assembly

Automated yeast assembly has three steps (i) DNA sample preparation, (ii) yeast transformation, (iii) transformation plating.

In step (i), a liquid handler distributes 15µL competent yeast cells + assembly vector (this is obtained by adding 5fmol/sample in less than 0.5µl/sample of assembly vector to the miniaturized competent cell master mix) into a 96 or 384 well plate. Insert DNA sample preparation can be performed in several ways depending on the application but in general, a traditional liquid handler or an acoustic liquid handler (Opentrons FLEX and Echo550 in this study) are programmed to dispense the desired insert segments on a 96 or 384-well plate. Typically, each well will contain 10-16 fmol per insert segment, about twice the amount needed for assembly.

In step (ii) the liquid handler is programmed to transfer 5-8 fmol of each DNA sample (8 fmol each insert fragment in less than a total of 2.5 µL) to the plate with competent yeast plus vector and to mix the components. The transformation plate is incubated on a heat block/thermocycler (Opentrons thermocycler module, Queens, NY, or INHECO 384 well cycler, Martinsried, Germany) with the appropriate adapter for the plate type. The transformation reaction is incubated at 30°C for 10-45 minutes and then at 42°C for 30 minutes.

In the final step (iii), the liquid handler adds 15µL of a solution consisting of 5% PEG-3350, 5mM CaCl_2_ into each well, aspirates 15µL, and for each sample, pipettes 2.5µL spots in a straight line spaced 2.6 mm apart on a Nunc Omnitray containing appropriate SC–Leu agar medium. The 48 samples are separated by 20mm horizontally, and 9mm vertically on the OmniTray, facilitating use of a standard vertical 8-channel pipette on the Opentrons Flex.

Resulting colonies are picked (either by hand or with a colony picker like the RapidPick SP (Hudson Robotics, Springfield NJ) into 100µL of sterile water, after which 30µL are transferred into a 384 well Labcyte Echo plate, and the Labcyte Echo robot is used to single colony purify (see **Supplementary text** for a description of the dispensing protocol).

### Oligo Guided Assembly Experiments

The built architectures (Fig. 6) were designed using a greedy algorithm that chose the next architecture based on the highest difference from all previously selected architectures, starting from a base architecture. The difference metric between architectures compares the location of each sub-assembly (so that the difference is higher if the subassembly is in a different location) and the orientation of each subassembly (so that the difference is higher if the subassembly is pointing in opposite direction, regardless of location).

For each architecture, the appropriate linker groups in sizes of either 300bp or 132bp were designed. For both sizes 16bp on each side were common primer binding sites to each linker group for specific amplification. For both sizes, the rest of the linker bases were split evenly to create overlaps with the right side of the sub-assemblies it is linking to; except that, for the left-most linker which in the middle has the 22bp Leu3 UAS, and overlaps with the SURE vector.

All oligonucleotide progenitors for the linkers were ordered in a pool (Twist Biosciences, Carlsbad, CA). The pool was resuspended at 0.5ng/µL. Each PCR reaction consisted of 2.4µL of the pool, 7.5µL of 2X Q5 polymerase (New England Biolabs, Ipswitch, MA), and 2.6µL of sterile water, and 2.5µL each of the appropriate 10µM specific primers. All reactions were set in a 96-well plate by distributing a master mix without primers and then using a liquid handler to add the primers. The reactions were set directly on the in-deck thermocycler, then the robot added a lid and initiated the PCR. Then the automated transformation was carried as above with the difference that all the YAV8 segments that were common to the assembly were added to the competent yeast in bulk. Colonies phenotypes were checked by plating to SC–Leu and SC–8 as described above for YAV8 colonies.

## Acknowledgments

We thank Henri Berger, Leslie A. Mitchell, and Jon Laurent for early work developing progenitors to the YAV8 system. This work was supported in part by grants R01HG012743 from the NIH/NHGRI to J.D.B. and U24HG011735 to Mark Adams, PI.

