## Supplement for "Assemblatron: An Automated Workflow for High-Throughput Assembly of Big-DNA Libraries"

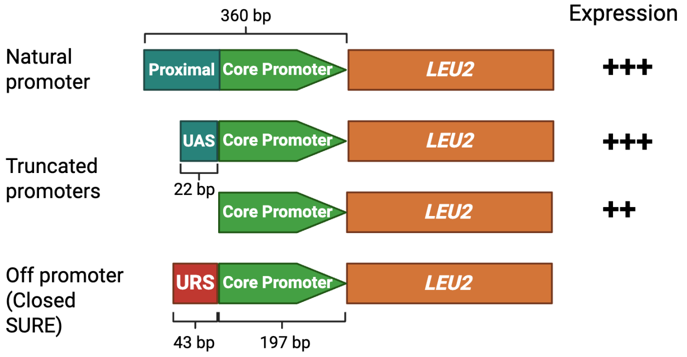


**Supplemental Figure 1. SURE vector control experiments.** The structures of four *LEU2* gene configurations is shown, along with the relevant expression phenotype, namely strength of growth on SC–Leu plates. Both truncated promoter configurations are functional in the context of SURE vector + inserts; see text for detail.

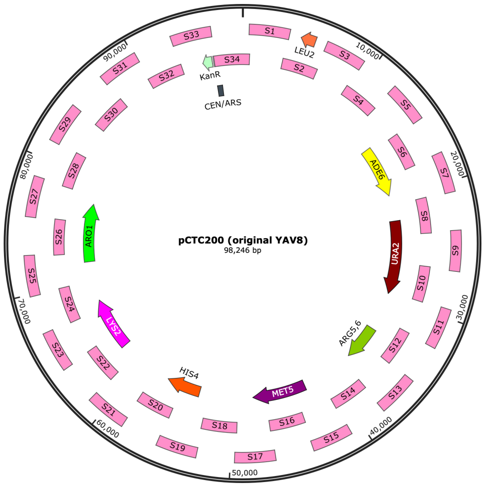


**Supplemental Figure 2**. YAV8 vector pCTC200 showing the 8 auxotrophic genes used as phenotypic indicators of proper yeast assembly, and 34 segments that can be used in its assembly. Mixing the 34 fragments and transforming strain yHB035 results in >95% correct assemblies as assayed by selection on SC–Leu plates followed by replica plating to SC–8 plates.

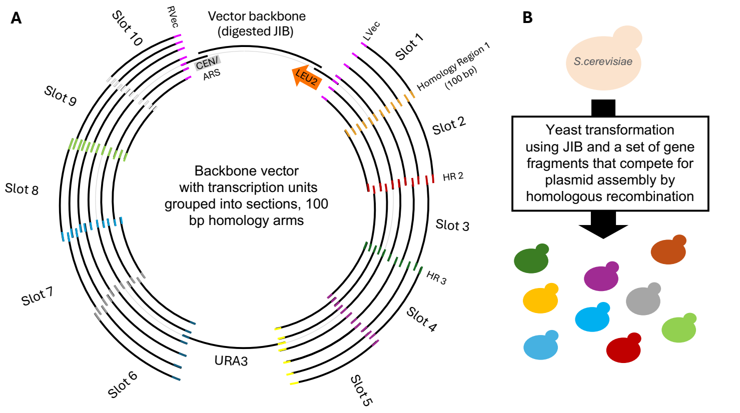


**Supplemental Figure 3.** Use of a JIB vector to produce a large library of biosynthetic pathways or other assemblies. 10 “slots” are shown. For pathway assembly the slots could harbor transcription units (TUs) driven by distinct promoters, orthologs from distinct organisms, or could be varied in other ways to generate diverse libraries that can be selected upon.

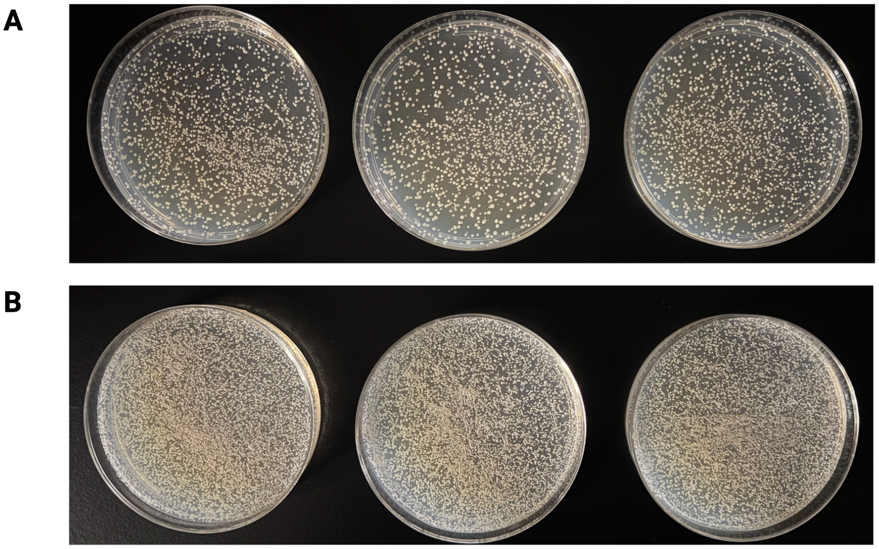


**Supplemental Figure 4. Transformation of a plasmid in a 96-well plate using the traditional competent cell preparation and our miniaturized preparation.** Yeast were made competent with the traditional or our improved methods (see **Methods** section). **A.** 30 µL of the traditional mix were transformed with 6.3 ng of pRS415 and plated in SC–Leu, obtaining a transformation efficiency of $10^5$ CFU/µg. **B.** 30 µL of the miniaturized mix were transformed with 6.3 ng of pRS415 and plated in SC–Leu, obtaining substantially more colonies.

**Supplemental Table 1 Plasmids.**

| Plasmid | Description |
|---|---|
| pAVG030 | JIB vector, LEU2 CEN |
| pAVG054 | JIB+SURE vector, LEU2 CEN |
| pAVG052 | SURE vector, LEU2 CEN |
| pAVG044 | "Assembled" JIB vector from pAVG030 |
| pAVG058 | SURE vector plus YAV8 inserts |
| pAVG059 | JIB vector plus YAV8 inserts |
| pKPR125 | JIB vector, LEU2 CEN, LVec_RVec |
| pJWM29 | (JIB for mSwAP-IN) |
| pKML70 | JIB_Rvec_Lvec in pUC19 |
| pKML72 | JIB+SURE in pUC19 |

**Supplemental Table 2. YAV8 gene parts**

| Gene | ORF size | intron | Gene coordinates | Genomic deletion coordinates |
|---|---|---|---|---|
| *ADE6* | 4077 | *ACT2* | chrVII:615965..611889 | chrVII:616495..611379 |
| *ARG56* | 2592 | *PHO85* | chrV: 295410..298001 | chrV:298511.. 299910 |
| *ARO1* | 4767 | *NSP1* | chrIV:704484..709250 | chrIV:703954..709750 |
| *HIS4* | 2400 | *OST5* | chrIII:68333..65934 | chrIII:68872.. 64549 |
| *LYS2* | 4179 | *SEC14* | chrII:473926..469748 | chrII:474216..469719 |
| *MET5* | 4329 | *BET1* | chrX:683285..678957 | chrX: 683786.. 678445 |
| *URA2* | 6645 | *SAC6* | chrX:172367..165723 | chrX:172494..165635 |
| *LEU2* | 1095 | none | chrIII:91324..92418 | chrIII:84794.. 92481 |

**Supplemental Table 3 yeast strains**

| | |
|---|---|
| BY4741 | *MAT**a** ura3Δ0 leu2Δ0 his3Δ1 met15Δ0* |
| BY4742 | *MATα ura3Δ0 leu2Δ0 his3Δ1 lys2Δ0* |
| YZY035 | *MATα ura3Δ0 leu2Δ0 lys2Δ0 (restored HIS3+ in BY4742)* |
| YZY501 | *MATα leu2Δ0 lys2Δ0 (restored URA3+ in YZY035)* |
| YHB35 | *MATα leu2Δ0 lys2Δ0 ade6Δ0 ura2Δ0 arg56Δ0* |
| YHB37 (YAVG1) | *MATα leu2Δ0 ade6Δ0 arg56 Δ0 aro1Δ0 his4Δ0 lys2Δ0 met5Δ0 ura2Δ0* |

**Supplemental Table 4 60 mer oligonucleotide sequences**

| Oligo# | Name | Sequence* |
|---|---|---|
| **1** | 1_Ura3 | gcctctcaccataacgtacgtagggataacagggtaatttgatttcggtttctttgaaat |
| **2** | 10_spHis5 | aaggaagtatatgaaagaagaacctcagtggcaaatcctaaccttttatatttctctaca |
| **3** | 10_Ura3 | caatgaagcacacaagtttgtttgcttttcgtgcatgatattaaatagcttggcagcaac |
| **4** | 11_spHis5 | cactgaggttcttctttcatatacttccttttaaaatcttgctaggatacagttctcaca |
| **5** | 11_Ura3 | gaaaagcaaacaaacttgtgtgcttcattggatgttcgtaccaccaaggaattactggag |
| **6** | 12_spHis5 | ctacccatggttgtttatgttcggatgtgatgtgagaactgtatcctagcaagattttaa |
| **7** | 12_Ura3 | acaaattttgggacctaatgcttcaactaactccagtaattccttggtggtacgaacatc |

| | | |
|---|---|---|
| **8** | 13_spHis5 | tcacatccgaacataaacaaccatgggtaggagggcttttgtagaaagaaatacgaacga |
| **9** | 13_Ura3 | ttagttgaagcattaggtcccaaaatttgtttactaaaaacacatgtggatatcttgact |
| **10** | 14_spHis5 | tccaaagcgatggcaacgctgattttcgtttcgttcgtatttctttctacaaaagccctc |
| **11** | 14_Ura3 | cggcttaactgtgccctccatggaaaaatcagtcaagatatccacatgtgtttttagtaa |
| **12** | 15_spHis5 | aacgaaaatcagcgttgccatcgctttggacaaagctcccttacctgaagagtcgaattt |
| **13** | 15_Ura3 | gatttttccatggagggcacagttaagccgctaaaggcattatccgccaagtacaatttt |
| **14** | 16_spHis5 | gcatgcttggaagttataagttcatcaataaaattcgactcttcaggtaagggagctttg |
| **15** | 16_Ura3 | gtcagcaaattttctgtcttcgaagagtaaaaaattgtacttggcggataatgcctttag |
| **16** | 17_spHis5 | tattgatgaacttataacttccaagcatgcaaaccaaaagggagaacaagtaatccaagt |
| **17** | 17_Ura3 | ttactcttcgaagacagaaaatttgctgacattggtaatacagtcaaattgcagtactct |
| **18** | 18_spHis5 | atgtgatccaagaatccaattcccgtgtctacttggattacttgttctcccttttggttt |
| **19** | 18_Ura3 | tgcccattctgctattctgtatacacccgcagagtactgcaatttgactgtattaccaat |
| **20** | 19_spHis5 | agacacgggaattggattcttggatcacatgtatcatgcactggctaaacatgcaggctg |
| **21** | 19_Ura3 | gcgggtgtatacagaatagcagaatgggcagacattacgaatgcacacggtgtggtgggc |
| **22** | 2_spacer | atttaattatatcagttattaccctgcggtgtgaaataccgccgatgctagcaaggcgaa |
| **23** | 2_Ura3 | cttctgttcggagattaccgaatcaaaaaaatttcaaagaaaccgaaatcaaattaccct |
| **24** | 20_spHis5 | aaatcacctcttgagtaaagtcgtaagctccagcctgcatgtttagccagtgcatgatac |
| **25** | 20_Ura3 | cgcctgcttcaaaccgctaacaatacctgggcccaccacaccgtgtgcattcgtaatgtc |
| **26** | 21_spHis5 | gagcttacgactttactcaagaggtgatttaatcatcgatgatcatcacactgcagaaga |
| **27** | 21_Ura3 | ccaggtattgttagcggtttgaagcaggcggcagaagaagtaacaaaggaacctagaggc |
| **28** | 22_spHis5 | ttgaatgcaataccaagtgcaatagcagtatcttctgcagtgtgatgatcatcgatgatt |
| **29** | 22_Ura3 | cttgcatgacaattctgctaacatcaaaaggcctctaggttcctttgttacttcttctgc |
| **30** | 23_spHis5 | tactgctattgcacttggtattgcattcaagcaggctatgggtaactttgccggcgttaa |
| **31** | 23_Ura3 | cttttgatgttagcagaattgtcatgcaagggctccctatctactggagaatatactaag |
| **32** | 24_spHis5 | tcaagtggacaataagcatgtccaaatcttttaacgccggcaaagttacccatagcctgc |
| **33** | 24_Ura3 | tttgtcgctcttcgcaatgtcaacagtacccttagtatattctccagtagatagggagcc |
| **34** | 25_spHis5 | aagatttggacatgcttattgtccacttgacgaagccctttctagaagcgtagttgactt |
| **35** | 25_Ura3 | ggtactgttgacattgcgaagagcgacaaagattttgttatcggctttattgctcaaaga |
| **36** | 26_spHis5 | aaatcgataacagcatagggccgtcccgacaagtcaactacgcttctagaaagggcttcg |
| **37** | 26_Ura3 | atcgtaaccttcatctcttccacccatgtctctttgagcaataaagccgataacaaaatc |
| **38** | 27_spHis5 | gtcgggacggccctatgctgttatcgatttgggattaaagcgtgaaaaggttggggaatt |
| **39** | 27_Ura3 | gacatgggtggaagagatgaaggttacgattggttgattatgacacccggtgtgggttta |
| **40** | 28_spHis5 | tatagtaagtgagggatcatttcacaggacaattccccaaccttttcacgctttaatccc |
| **41** | 28_Ura3 | ctgttgacccaatgcgtctcccttgtcatctaaacccacaccgggtgtcataatcaacca |
| **42** | 29_spHis5 | gtcctgtgaaatgatccctcacttactatattccttttcggtagcagctggaattacttt |
| **43** | 29_Ura3 | gatgacaagggagacgcattgggtcaacagtatagaaccgtggatgatgtggtctctaca |
| **44** | 3_spacer | ctcatcgaccgcagcctagctatacgatcgttcgccttgctagcatcggcggtatttcac |
| **45** | 3_spHis5 | tccttgacagtcttgacgtgcgcagctcaggggcatgatgtgactgtcgcccgtacattt |
| **46** | 3_Ura3 | ttttttgattcggtaatctccgaacagaaggaagaacgaaggaaggagcacagacttaga |

| | | |
|---|---|---|
| **47** | 30_spHis5 | tcattactaccatataagcaggtaacatgcaaagtaattccagctgctaccgaaaaggaa |
| **48** | 30_Ura3 | tcctcttccaacaataataatgtcagatcctgtagagaccacatcatccacggttctata |
| **49** | 31_spHis5 | gcatgttacctgcttatatggtagtaatgaccatcatcgtgctgaaagcgcttttaaatc |
| **50** | 31_Ura3 | ggatctgacattattattgttggaagaggactatttgcaaagggaagggatgctaaggta |
| **51** | 32_spHis5 | ctagtagccgcgcgcatggcaacagccagagatttaaaagcgctttcagcacgatgatgg |
| **52** | 32_Ura3 | ccagcctgcttttctgtaacgttcaccctctaccttagcatcccttccctttgcaaatag |
| **53** | 33_spHis5 | tctggctgttgccatgcgcgcggctactagtcttactggaagttctgaagtcccaagcac |
| **54** | 33_Ura3 | gagggtgaacgttacagaaaagcaggctgggaagcatatttgagaagatgcggccagcaa |
| **55** | 34_spHis5 | attgtcagtactctttacaacactcccttcgtgcttgggacttcagaacttccagtaaga |
| **56** | 34_Ura3 | tgcatttacttataatacagtttttttagttttgctggccgcatcttctcaaatatgcttc |
| **57** | 35_spHis5 | gaagggagtgttgtaaagagtactgacaataaaaagattcttgttttcaagaacttgtca |
| **58** | 35_Ura3 | aactaaaaaactgtattataagtaaatgcatgtatactaaactcacaaattagagcttca |
| **59** | 36_spHis5 | aacaactacaatataaaaaaactatacaaatgacaagttcttgaaaacaagaatcttttt |
| **60** | 36_Ura3 | accgcagggtaataactgatataattaaattgaagctctaatttgtgagtttagtataca |
| **61** | 37_spHis5 | tttgtatagtttttttatattgtagttgttctattttaatcaaatgttagcgtgatttat |
| **62** | 38_spHis5 | tgggcagatgatgtcgaggcgaaaaaaaatataaatcacgctaacatttgattaaaatag |
| **63** | 39_spHis5 | atttttttcgcctcgacatcatctgcccagatgcgaagttaagtgcgcagaaagtaata |
| **64** | 4_spacer | cgatcgtatagctaggctgcggtcgatgagcttaggagacatggaggcccagaatacccg |
| **65** | 4_spHis5 | caaatgattatacatggggatgtatgggctaaatgtacgggcgacagtcacatcatgccc |
| **66** | 4_Ura3 | ttcaacactacatatgcgtatatataccaatctaagtctgtgctccttccttcgttcttc |
| **67** | 40_spHis5 | gaccagcattcacatacgattgacgcatgatattactttctgcgcacttaacttcgcatc |
| **68** | 41_spHis5 | tcatgcgtcaatcgtatgtgaatgctggtcgctatactgattaccctgttatccctagat |
| **69** | 42_spHis5 | tacgaggcgcgtgtaagttacaggcaagcgatctagggataacagggtaatcagtatagc |
| **70** | 5_spacer | ctgagctgcgcacgtcaagactgtcaaggacgggtattctgggcctccatgtctcctaag |
| **71** | 5_spHis5 | agcccatacatccccatgtataatcatttgcatccatacattttgatggccgcacggcgc |
| **72** | 5_Ura3 | ttggtatatatacgcatatgtagtgttgaagaaacatgaaattgcccagtattcttaacc |
| **73** | 6_spHis5 | tctgcagcgaggagccgtaatttttgcttcgcgccgtgcggccatcaaaatgtatggatg |
| **74** | 6_Ura3 | tttcctgcaggtttttgttctgtgcagttgggttaagaatactgggcaatttcatgtttc |
| **75** | 7_spHis5 | gaagcaaaaattacggctcctcgctgcagacctgcgagcagggaaacgctcccctcacag |
| **76** | 7_Ura3 | caactgcacagaacaaaaacctgcaggaaacgaagataaatcatgtcgaaagctacatat |
| **77** | 8_spHis5 | ggggcgcggcgtggggacaattcaacgcgtctgtgaggggagcgtttccctgctcgcagg |
| **78** | 8_Ura3 | aggactaggatgagtagcagcacgttccttatatgtagctttcgacatgatttatcttcg |
| **79** | 9_spHis5 | acgcgttgaattgtccccacgccgcgcccctgtagagaaatataaaaggttaggatttgc |
| **80** | 9_Ura3 | aaggaacgtgctgctactcatcctagtcctgttgctgccaagctatttaatatcatgcac |
| **81** | SURE_ad-apter_UAS | gttatccctacgtacgttatggtgagaggccggaaccggcttttcatatagaatagagaagcgttcatga |

*All oligos are 60mers except the last one which is a 70mer

**Supplemental text**

**Single Colony Purification with Echo**

The Echo protocol uses a Nunc Omnitray divided into 192 columns and 128 rows, forming 24576 destination points. Each source well (which represents one colony) is dispensed in an area of 2 columns by 49 rows, the Echo shots are coordinated so from top to bottom the frequency is decreasing and the spacing between shots is increasing. At the highest density 50 nL are shot into a 2x2 points area, at the lowest density 2.5 nL are shot in a 2x5 points area (this is effectively one 2.5 nL shot within the 2x5 area). One point is a linear travel distance of about 0.57 mm, the smallest distance we could program on the Echo's stepper motor. This creates a dilution which serves to isolate single colonies. Each area is separated horizontally and vertically from the next by 6 columns/rows, which gives enough space for colonies to grow without touching each other. In this configuration it is possible to single colony purify 48 colonies on an OmniTray.
